# Novelty and anxiety differentially shape hippocampal assembly reactivation and sequence replay during REM sleep

**DOI:** 10.1101/2024.09.05.611474

**Authors:** Jisoo Choi, James Carmichael, Sylvain Williams, Guillaume Etter

## Abstract

Mammalian sleep alternates between rapid eye movement (REM) and non-REM (NREM) phases, each with distinct patterns of neural activity. The replay of waking experience during NREM sleep is a well-established substrate for memory consolidation, but whether comparable reactivation occurs during REM sleep has remained unclear, owing to the sparse firing that characterizes this state and the lack of robust methods for detecting replay within it. Here we combined calcium imaging with electrophysiology to monitor large populations of hippocampal CA1 neurons in mice and applied complementary analytical approaches to test for neural reactivation during REM sleep. We found that structured waking activity is re-expressed during REM sleep across a range of temporal organizations, from the reactivation of co-active cell assemblies to temporally ordered sequential replay. This reactivation was selectively shaped by the behavioral context of prior experience. Novel experience enhanced sequential replay and refined assembly composition, selectively reducing the participation of weakly contributing neurons; anxiogenic experience, by contrast, increased the frequency of assembly reactivation and reinforced the recruitment of strongly contributing neurons. REM sleep thus emerges as an active stage of memory processing that reorganizes recent experience according to its behavioral significance, engaging distinct operations for novel and for emotionally salient events.

**Significance statement:** Replay of waking neural activity during non-REM (NREM) sleep is strongly linked to memory consolidation, but technical and computational obstacles have left the role of REM sleep largely unresolved. Combining large-scale hippocampal recordings across wakefulness and REM sleep with complementary computational approaches, we show that REM sleep robustly reactivates activity patterns formed during awake experience. This reactivation is shaped by the behavioral context of the preceding experience, with novelty and anxiety engaging distinct forms. These findings identify REM sleep as an active brain state that engages distinct operations for novel and for emotionally salient experiences, offering new insight into how REM sleep may contribute to memory consolidation.

## Introduction

Sleep-dependent memory consolidation has been studied predominantly during NREM sleep, which has a well-established causal role in this process^1–4^. Short hippocampal events called sharp-wave ripples (SWRs), observed during NREM sleep, coincide with reactivations of place cell sequences that reflect previous awake experiences^4–9^. Disrupting or prolonging SWRs and their associated replay events detrimentally affects or strengthens spatial memory, respectively, highlighting their critical role in memory consolidation^1–3,9,10^. Most current models of offline hippocampal processing derive from this work.

Considerably less is known about hippocampal activity during REM sleep. Rather than discrete SWR events, REM sleep is characterized by sustained hippocampal theta oscillations similar to those observed during wakefulness^11,12^, and neuronal firing during this state is sparse and less frequent. During wakefulness, theta oscillations organize place cell firing into structured sequences^13^, raising the possibility that activity patterns formed during theta-associated exploration are re-expressed during the theta-dominated state of REM sleep. However, both the existence of such re-expression and its potential contribution to memory consolidation remain unclear, with the available evidence limited and inconsistent. Early studies reported that reactivation of awake hippocampal activity during REM sleep was either absent^14^ or observed occasionally^15^. Other research showed that place cells active during wakefulness resumed firing in REM sleep and displayed experience-dependent phase-locking to theta oscillations^16^. A recent large-scale electrophysiological study confirmed strong reactivation of place cells during SWRs in NREM sleep but found only weak reactivation during REM-associated theta oscillations^17^. The causal evidence is similarly limited: optogenetic suppression of theta oscillations during REM sleep impairs contextual fear and novel object spatial memory^11^, and optogenetic manipulation of ensembles of adult-born dentate gyrus neurons during REM sleep impairs contextual fear memory^18^. Together, these findings establish a causal role for REM sleep in memory consolidation, but the structured neural activity patterns that could mediate this role remain unidentified. This gap is attributable in part to the sparse neuronal activity characteristic of REM sleep, to methodological limitations of conventional single-unit electrophysiological recordings, and to the use of only a single computational approach for detecting reactivation in most studies. In addition, existing studies of REM reactivation have been largely restricted to well-trained animals in highly familiar or similar environments^15^, limiting our understanding of how recent experience shapes reactivation. This gap is especially significant because REM sleep has been implicated in the processing of emotionally salient experiences^19–21^, yet the neural activity patterns that support this function remain unknown.

Here we combined in vivo calcium imaging with electrophysiological recordings to monitor large populations of CA1 neurons, including hundreds of place cells per animal, across wakefulness and REM sleep, before and after a behavioral task, an approach that holds neuronal identity fixed across brain states and offsets the sparse firing of REM sleep. Because the temporal organization of any reactivated structure was unknown, we applied three analyses imposing progressively stronger assumptions about temporal order, from co-activity of cell assemblies through fixed-order sequences to decodable spatial trajectories, and examined how reactivation at each level is shaped by novel and by anxiogenic experience. We find that REM sleep robustly re-expresses waking activity, from co-active assemblies to ordered sequences, and that novelty and anxiogenic experience engage dissociable operations on these two forms of reactivation.

## Results

### Large-scale recording of hippocampal activity across wake and REM sleep

Whether awake hippocampal activity is reactivated as recurring neural activity patterns during REM sleep remains an open question. To address this, we combined calcium imaging with electrophysiology to simultaneously record large neuronal populations(>1,000 cells per animal), substantially improving our ability to characterize neural activity patterns formed during awake experience and to track their reactivation across REM sleep (Fig. 1a,b). Calcium imaging was performed with open-source miniscopes^22^, and single-neuron signals were extracted using CNMFe^23^ (Fig. 1c). Each session comprised 2 h of pre-task sleep in the home cage, a 10 min behavioral task, and 2 h of post-task sleep. Animals were recorded first on the linear track across multiple days, then on the anxiety track, which was walled along only half of its length (Fig. 1d; see Methods). REM episodes were manually identified during sleep sessions and subsequently verified offline based on the theta/delta ratio and EMG amplitude (Fig. 1e,f). REM epochs showed a markedly higher theta/delta ratio than NREM epochs (REM: 5.49 ± 0.43; NREM: 0.84 ± 0.04; t(166) = 10.38, p < 0.001; Fig. 1f). REM sleep (Pre-REM: 4.35 ± 1.71min; Post-REM: 5.23 ± 1.72 min) occupied the smallest fraction of each session (7.07 ± 0.45%), compared with wake (31.55 ± 1.46%) and NREM (60.50 ± 1.20%; RM-ANOVA, F(2,24) = 387.06, p < 0.0001, η² = 0.970; Fig. 1g).

**Figure 1.**
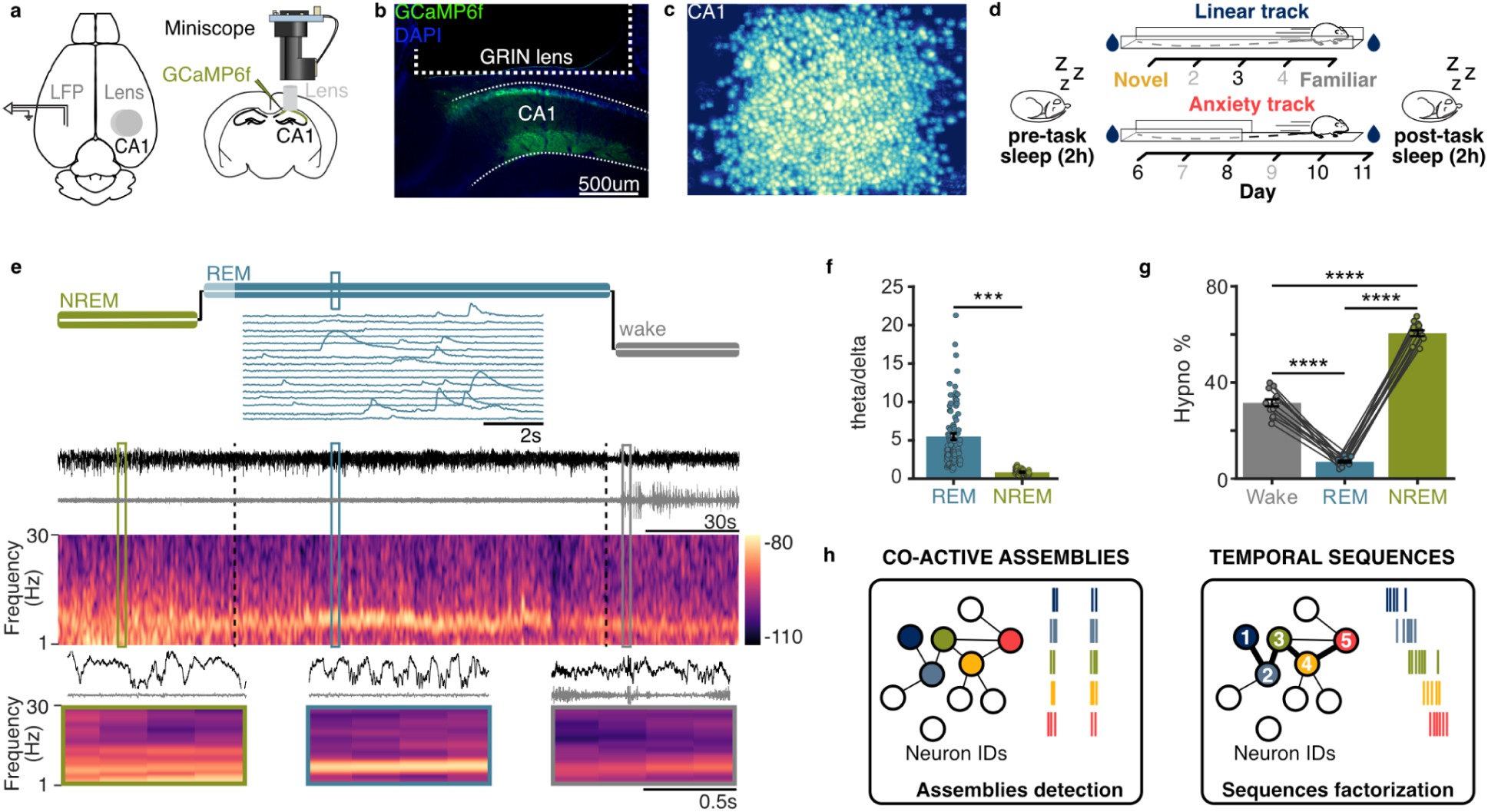
Recording hippocampal activity across wake and REM sleep. **a**, CA1 neurons were transfected with GCaMP6f, and a GRIN lens was implanted above the pyramidal cell layer for imaging; tungsten wires were implanted contralaterally for local field potential (LFP) recording. **b**, Histology showing GCaMP6f expression in the CA1 pyramidal layer and the corresponding lens position. **c**, Individual CA1 neurons extracted from calcium imaging videos. **d**, Each recording session included sleep before and after a behavioral task. Animals were first recorded on the linear track and subsequently on a partially walled anxiety track across days. **e**, REM sleep calcium recordings were manually acquired when theta oscillations occurred together with minimal EMG activity (inset: raw Ca²⁺ transients from 16 example cells). **f**, Sleep states were rescored offline based on the theta/delta ratio and root-mean-square EMG amplitude. REM epochs showed a significantly higher theta/delta ratio than NREM. **g**, Percentage of session time spent in wake, REM, and NREM. **h**, Hypothesized forms of hippocampal reactivation during REM sleep, spanning low temporal organization (co-active assemblies) to highly structured temporal sequences. Data expressed as mean ± SEM. ***, *p* < 0.001; ****, *p* < 0.0001. Test used in **f**: independent t test; **g**: Repeated measures ANOVA.

We reasoned that preconfigured internal dynamics^24^ could shape hippocampal activity during REM sleep, and that any resulting reactivation need not take a single form: it could range from loosely structured co-active assemblies to precisely ordered temporal sequences, which are not mutually exclusive (Fig. 1h). Using complementary analytical approaches (see Methods), we therefore asked whether structure formed during awake experience is reactivated during subsequent REM sleep, and how such reactivation is shaped by exposure to novel, familiar, or anxiogenic contexts.

### Detecting co-active cell assembly and temporal sequence reactivation in REM sleep

We first asked whether hippocampal patterns expressed during the mice’s first exposure to a linear track are reactivated during subsequent REM sleep (Fig. 2a). We focused on place cells (Fig. 2b–d), defined as the neurons with significant spatial tuning and the highest spatial information content (top 256 cells per mouse; see Methods), ensuring robust spatial encoding. Restricting the analysis in this way increased the likelihood of detecting meaningful reactivations, even within relatively small neuronal populations.

**Figure 2.**
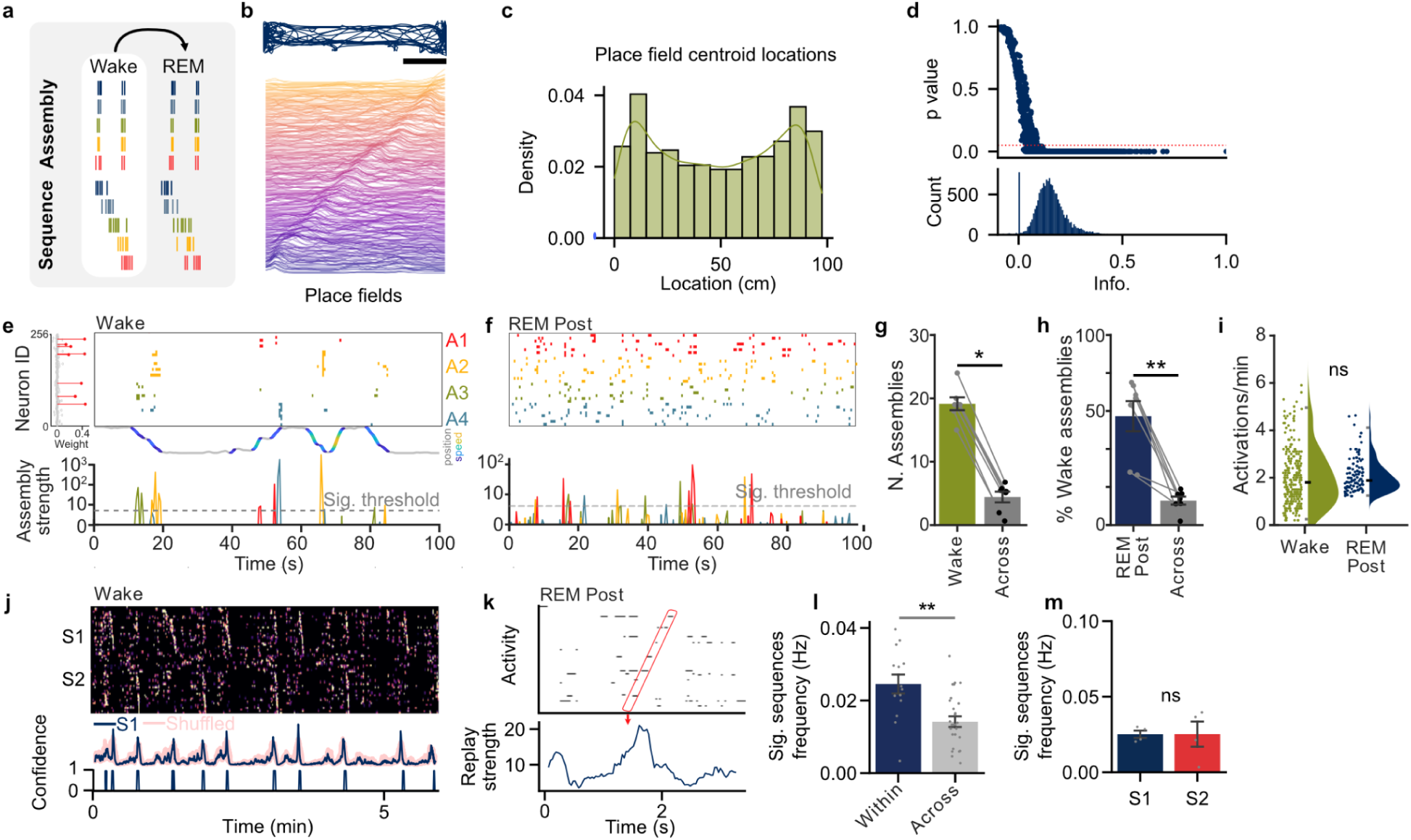
Reactivations of awake activity patterns in CA1 during subsequent REM sleep. **a,** We evaluated whether neural activity patterns from awake experience were recapitulated during subsequent REM sleep. **b**, Example of mouse trajectory on a linear track (top) and corresponding place fields of recorded CA1 neurons, sorted by centroid location (bottom). Each place field is normalized by its peak activity. **c**, Distribution of place field centroids on the linear track. **d**, Spatial information and significance for all recorded neurons (bottom) and associated distribution (top). **e**, Example assemblies detected in awake activity, with neuron weights shown for one assembly (A1, red). **f**, Examples of significant reactivations of wake assembly patterns during post-task REM sleep. The number of detected assemblies in wake (**g**) and post-task REM sleep (**h**) exceeded chance levels derived from across session surrogates. **i**, Assembly activation rates in wake, and reactivation rates of wake assemblies in post-task REM, both exceeded chance levels estimated from shuffled surrogates (gray). **j**, Example sequential activity for each running direction (S1, S2) during linear-track exploration (top), tested against shuffled surrogates (middle), with the resulting sequence confidence (bottom). **k**, Example sequence detected during subsequent REM sleep, with replay strength plotted over time. **l**, Expression of sequences in REM sleep exceeded chance (across-sessions surrogates). **m**, Frequency of sequential activity during REM sleep did not differ between running directions. Data expressed as mean ± SEM. ns, not significant; *, *p* < 0.05; **, *p* < 0.01. Test used in **g**: wilcoxon sign rank test; **h**: paired t test; **i**: independent t test; **l**: Kruskal-Wallis test.

We began with the simplest assumption about temporal structure: that neurons firing together during wakefulness might do so again during REM sleep, irrespective of their order. We therefore tested whether a subset of these place cells formed assemblies of co-active neurons (Fig. 2e). Assemblies were identified by applying principal component analysis to correlated wake activity (0.5s bins), retaining components whose eigenvalues exceeded the 99th percentile of the shuffled data. Independent component analysis was then applied to quantify each neuron’s contribution (weight) to each assembly^25^ (see Fig. S1 for visualized methods). Wake-derived assembly templates were then applied to REM data from the home-cage recordings (Fig. 2f), and time points at which assembly co-activity exceeded the significance threshold (99th percentile of shuffled data) were counted as significant reactivations. During linear-track running, we detected 19.14 ± 2.67 assemblies per session (n = 4 mice; Fig. 2g), significantly higher than the chance level derived from alternative session surrogates (4.43 ± 2.24; Wilcoxon signed-rank, w = 28, p = 0.016; see Methods). Of the detected assemblies in wake, 57.24% (± 7.90) displayed reactivations during subsequent REM sleep, exceeding the chance detection levels (12.79 +/-2.10%, REM shuffle: 2.50 ± 1.09%, F(1,6) = 157.67, p < 0.0001, Fig 2h). The rate of wake activation (1.99 ± 0.08 min^−1^) and subsequent REM reactivations across assemblies (2.06 ± 0.06 min^−1^) did not differ (paired t test, t(388) = -0.68, p = 0.50, Cohen’s d = -0.071; Fig. 2i). Thus, co-active assemblies formed during awake exploration are re-expressed during subsequent REM sleep at rates comparable to those observed during wakefulness.

We next asked whether REM activity also carried finer temporal structure, in the form of decodable spatial trajectories. We applied Bayesian decoding, the standard method for replay detection during SWRs in electrophysiological recordings, to awake-trajectory events (Fig. S2). Given the known limitations of Bayesian decoding using calcium imaging with lower temporal resolution, we validated the approach by comparing the number of decoded replay events per session against a null distribution generated by shuffling across sessions originating from different subjects. Briefly, in this control condition, Bayesian posterior probabilities were computed using priors (tuning curves) originating from distinct mice. The observed number did not exceed this null distribution, indicating that decoded trajectories during REM sleep were not reactivated above chance (Fig. S2g) within this time frame. We therefore did not pursue Bayesian decoding further.

We next asked whether REM activity carried finer temporal structure, in the form of ordered sequences: recurring motifs in which the same neurons fire in the same relative order, rather than decodable spatial trajectories. Using sequence factorization^26^, which identifies recurring temporal firing motifs without reference to the animal’s position, we extracted sequence templates from awake activity and applied the same templates to REM data, testing against shuffled surrogates using previously optimized parameters (see Methods; Fig. S3a-e). Two temporally structured sequences were extracted during exploration, corresponding to the two running directions (Fig. 2j), as expected given the directional specificity of place cells on a linear track. Importantly, the same sequences were significantly re-expressed during REM sleep following this novel experience (Fig. 2k,l), indicating that awake firing motifs recur in REM with their temporal order preserved.

Replay of these sequences was more frequent within sessions (0.0245 ± 0.003 Hz) than across sessions and subjects (0.0141 ± 0.001 Hz; independent t test, t(21.37) = 3.45, p = 0.0023, Cohen’s d = 1.223; Fig. 2l), with comparable replay frequencies for both running directions (Kruskal-Wallis test, H = 0.0833, p = 0.7728; Fig. 2m). The highest number of replay events was obtained when no expansion or compression factor was applied to the running template (Fig. S3e). Together, these results show that REM sleep contains re-expression of awake firing motifs with their temporal order preserved, even though these events were not detectable as decoded spatial trajectories.

### Novelty increases the frequency of ordered sequential replay and tunes assembly reactivation during REM sleep

Having established that awake activity is re-expressed during REM sleep, we asked how this re-expression is shaped by novelty. We compared REM sleep following initial exposure to a novel linear track (day 1) with REM sleep after five days of training on the same track (day 5; Fig. 3a). We first compared the reactivation of co-active assemblies, defined during track running, between novel and familiar sessions (Fig. 3b). The overall rate of assembly reactivation did not differ between conditions (novel: 2.24 ± 0.13/min; familiar: 1.96 ± 0.08/min; unpaired t-test t(59) = 1.96, p = 0.054, Cohen’s d = 0.50). We therefore asked whether novelty instead altered assembly composition. For each neuron we computed a participation change score, its probability of being active during assembly reactivations in post-task REM divided by the corresponding probability in pre-task REM (see Methods, Fig. S1n). A score of 1 indicates stable participation across pre- and post-task REM, whereas values below 1 indicate reduced participation during post-task REM. Assembly members were defined from the ICA-derived weights as neurons with z-scored weights exceeding 2 SD above the mean; all remaining neurons were classified as non-members. The participation change score for member-cells was lower from pre- to post-task REM sleep in the novel exposure (0.92 ± 0.06) compared to the familiar (1.14 ± 0.08; t test, t(129) = -2.27, p = 0.025, Cohen’s d = -0.40 Fig. 3c). Non-member cells participated far less in post-task reactivations following novel than familiar exposure (novel: 0.25 ± 0.02; familiar: 0.44 ± 0.03; unpaired t test, t(109) = −6.15, p < 0.0001, Cohen’s d = −1.07; Fig. 3d). Novel experience therefore refines cell assembly composition asymmetrically, shedding weakly contributing neurons during subsequent REM sleep while tuning the core membership.

**Figure 3.**
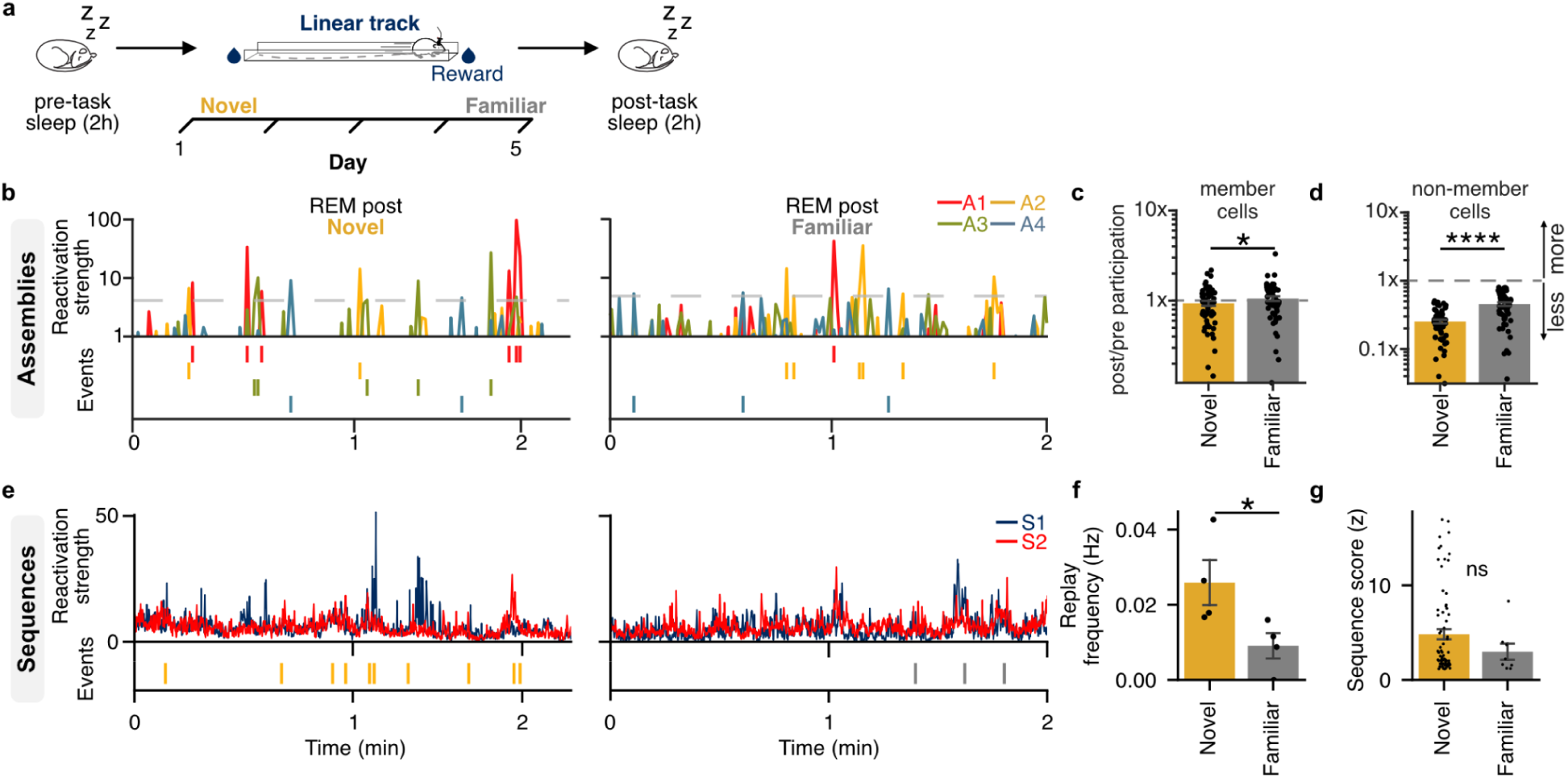
Novelty tunes assembly participation and increases the frequency of sequential replay. **a**, Mice explored the same linear track for five days; we compared REM hippocampal activity on day 1 (novel) with that on day 5 (familiar). **b**, Reactivation strength for representative assemblies (top) and the corresponding reactivation timestamps (bottom) following novel and familiar exposure. **c**, Participation of highly weighted ‘member’ cells was relatively similar between pre-and post-task REM session, with a lower overall participation following novel vs familiar exposure (score > 1 represents enhanced participation post-task). **d**, Participation of ‘non-member’ cells was reduced post-task, and significantly lower following novel compared to familiar exposure. **e**, Replay strength for both directional sequences (top) and the corresponding significant replay timestamps (bottom) following novel and familiar exposure. **f**, Sequences were replayed more frequently during REM sleep following novel than familiar exposure, whereas the sequence score of individual replay events did not differ (**g**). Data expressed as mean ± SEM. ns, not significant; *, *p* < 0.05; ****, *p* < 0.0001. Test used in **c/d/g**: independent t test; **f**: Repeated measures ANOVA.

We next asked whether novelty also affected temporal sequential replay (Fig. 3e). Sequences were replayed more frequently following novel than familiar exposure (novel: 0.026 ± 0.006 Hz; familiar: 0.009 ± 0.003 Hz; RM-ANOVA, F(1,3) = 33.84, p = 0.010, η² = 0.749, n = 4 mice; Fig. 3f), whereas the sequence score of individual replay events was unchanged across conditions (Fig. 3g; see Methods). These findings indicate that REM sleep processes novel experience through two complementary operations: it strengthens sequential replay by increasing frequency and refines assembly composition by pruning weakly participating neurons.

### Anxiogenic experience modulates assembly but not sequence reactivation in REM sleep

Given the proposed role of REM sleep in emotional memory processing^19,27–29^, we asked whether exposure to a naturally aversive elevated open space influences hippocampal reactivation during subsequent REM sleep. We built an “anxiety track”, an elevated linear maze in which one half was open (anxiogenic) and the other enclosed by walls (safe). Although mice spent more time in the enclosed portion, consistent with avoidance behavior, they were required to traverse the entire trajectory to receive a reward (Fig. 4a, Fig. S4).

**Figure 4.**
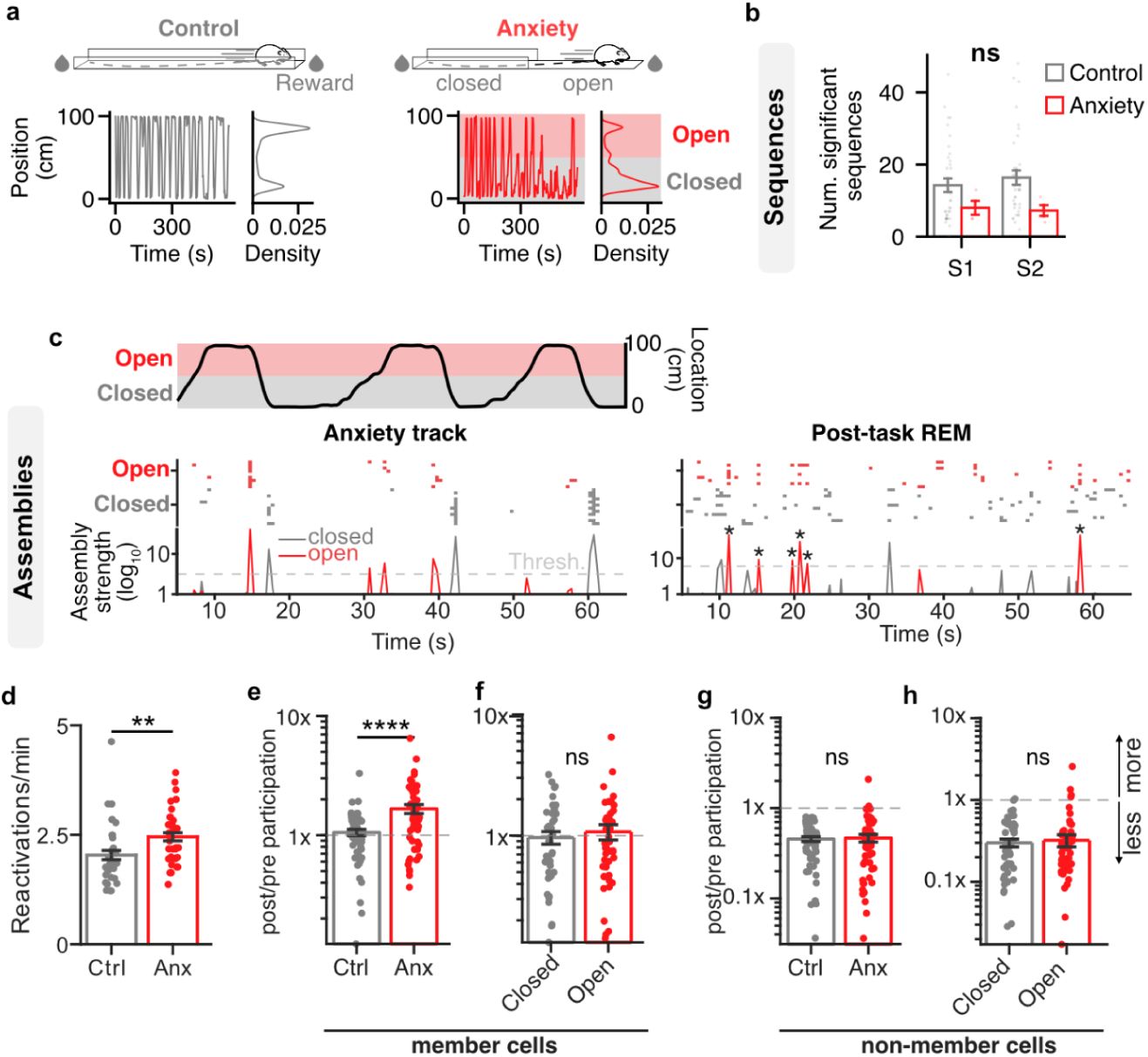
Anxiogenic experience increases the rate of assembly reactivation and the recruitment of strongly contributing cells in post-task REM. **a**, Trajectories and occupancy rate of the mouse in the enclosed control linear track and half enclosed anxiety track. Mice displayed avoidance, spending less time in the open portion of the anxiety track. **b**, Number of ordered sequences replayed during REM sleep following the anxiety track, relative to the linear track (control), separated by running direction. S1 and S2 denote direction-specific sequences; on the anxiety track these correspond to trajectories toward the open and closed arms, respectively. **c**, Example assemblies during awake exploration of the anxiety track (left) and their reactivation during subsequent REM sleep (right). The black trace shows the animal’s trajectory (red, open arm; grey, closed arm); the raster shows binarized activity of member neurons during locomotion; red and grey traces in the bottom panel show assembly strength for exemplar open-and closed-arm assemblies (stars, significant reactivations of an open-arm assembly). **d**, Exposure to the anxiety track increased the rate of offline assembly reactivation relative to the control linear track. **e**, Anxiogenic experience increased the participation of highly weighted ‘member’ cells relative to the control linear track, with no difference between open- and closed-arm-biased assemblies (**f**). **g**, Anxiogenic experience did not alter ‘non-member’ cell participation; **h**, non-member cell participation did not differ between open- and closed-arm assemblies. Data expressed as mean ± SEM. ns, not significant; **, *p* < 0.01; ****, *p* < 0.0001. Test used in **b**: Repeated measures ANOVA; **d-h**: independent t test.

Because the anxiety track has two directional components, toward and away from the open arm, we first asked whether ordered sequential replay was biased by direction. We found similar numbers of sequences were replayed toward and away from the open arm, and the overall rate of sequential replay did not differ between the anxiety track and the enclosed ‘control’ linear track (Fig. 4b). Sequence scores were also not different across control (1.23 ± 0.11) and anxiety (1.67 ± 0.19) conditions (RM-ANOVA, F_1,2_ = 2.2276, p = 0.1958; Fig. S4e). We then examined the impact of anxiogenic experience on co-active assemblies (Fig. 4c). Assembly reactivation was more frequent following the anxiety track (2.45 ± 0.09 min^−1^) than the control linear track (2.04 ± 0.11 min^−1^; unpaired t test, t(81) = −2.95, p = 0.0042, Cohen’s d = −0.64; Fig. 4d).

We next asked whether assembly composition was also altered by anxiogenic experience. Using the same participation change score as above, we found that member-cell participation was stronger following the anxiety track (1.66 ± 0.14) than the control linear track (1.06 ± 0.06; t(111) = −3.91, p < 0.001, Cohen’s d = −0.73; Fig. 4e), with no bias toward assemblies whose overall place tuning was biased to the open (1.56 ± 0.17) versus closed arm (1.50 ± 0.30) in anxiety track (t test, t(108) = 0.20, p = 0.84, Cohen’s d = 0.04; Fig. 4f). Interestingly, member-cell participation decreased as the mice became familiar with the anxiety track (‘Anx fam’) but increased again when the track configuration was altered by switching the open wall to the opposite direction (‘Anx Switch’, Fig. S4c). Non-member participation was unaffected by anxiogenic exposure, habituation to the anxiety track, or the subsequent track configuration switch (one-way ANOVA, F(3,245): 0.10 p = 0.96), and showed no bias toward the open (0.41 ± 0.05) versus closed arm (0.37 ± 0.04; t test, t(143) = 0.60, p = 0.550, Cohen’s d = 0.10; Fig. 4h, Fig. S4d). Anxiogenic experience therefore reshapes REM sleep assembly dynamics in a manner opposite to novelty: rather than pruning weakly engaged neurons, it directly modulates participation of the member cells in relation to behavioral relevance of the experience with the increased frequency of assembly reactivation.

### Organized neural activity in REM sleep precedes novel experience

Structured sequences that appear before the relevant waking experience, termed preplay, have been reported during NREM sleep^30–32^. Although preplay has been debated^33^, there is growing evidence that the hippocampus intrinsically generates structured patterns reflecting its developmental connectivity^34^; therefore, preconfigured patterns of activity must be considered alongside experience-driven replay. Whether preplay also occurs during REM sleep, however, has not been examined. We therefore evaluated two forms of structured activity in REM sleep: first, does REM sleep contain intrinsic assembly and/or sequential patterns before any task experience; and second, are these preconfigured patterns later expressed during wakefulness (Fig 5a)?

**Figure 5.**
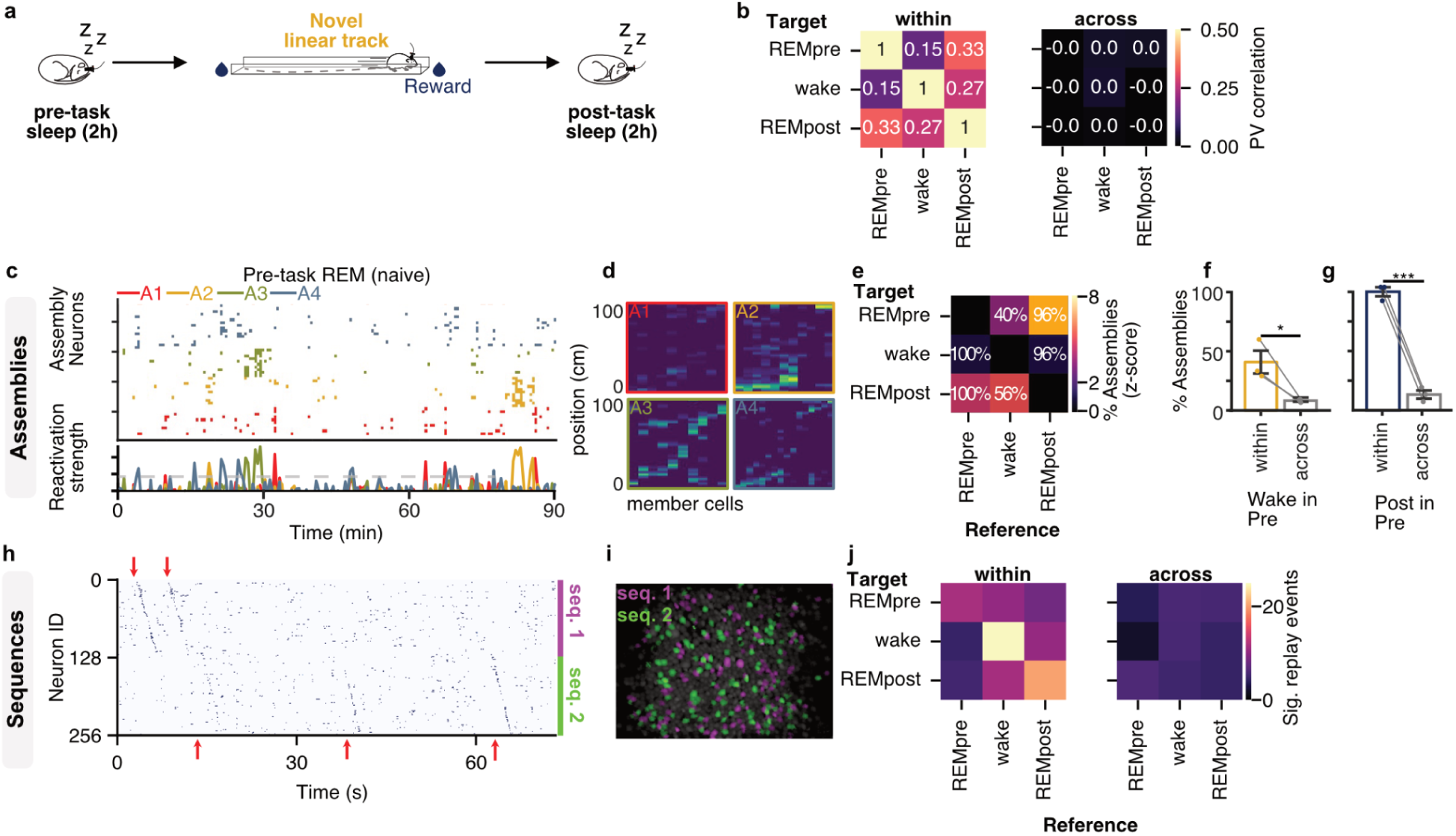
The internal structure of REM sleep activity precedes experience and is re-expressed during wakefulness. **a**, The day-1 track session was the animal’s first experience outside the home cage; with the preceding pre-task REM sleep representing a naïve state. **b**, Population vector (PV) correlation across pre-task REM, wakefulness, and post-task REM on day 1. Values represent the mean PV correlation for each comparison both within session (left) and using across session surrogates (right). **c**, Example preconfigured assemblies detected during REM sleep before the novel experience, showing co-activity of their constituent neurons. **d**, Spatial tuning maps on the subsequent novel track for the neurons composing the example pre-task REM assemblies. **e**, Percentage of significantly reactivated assemblies across conditions z-scored relative to other across subject controls (templates detected in the reference condition were applied to the activity in the target condition; text represents the raw percentages). **f**, The percentage of assemblies in wake that were expressed in naïve pre-task REM compared to across session surrogates. **g**, Percent of post-task REM assemblies expressed in naïve pre-task REM against across session surrogates. **h**, Preconfigured sequences detected during REM sleep before a novel experience in a representative mouse. **i**, Spatial footprints of the constituent neurons for sequences 1 and 2. **j**, Number of significant ordered sequences for each condition within subjects and across subjects (used as a control). Data expressed as mean ± SEM. ns, not significant; *, *p* < 0.05; ***, *p* < 0.001. Test used in **f/g/i**: paired t test.

Initially, we asked whether the identity of the active population, rather than its temporal structure, was preserved across states, by computing population vector (PV) correlations between states. Neurons active during REM sleep preceding a novel experience were most strongly correlated with those active during subsequent REM sleep (PV correlation: 0.33 ± 0.09; n = 4 mice), followed by the correlation between awake exploration and the following REM sleep episode (0.27 ± 0.013) and these values dropped to zero when PV correlation was computed across, rather than within mice (n = 4 mice; Fig. 5b).

To determine if awake assembly patterns were present in naïve pre-task REM sleep, we first applied assembly templates from the first track session to the pre-task REM data. We found that awake assemblies were active beyond the shuffle reactivation (‘r’) threshold (40.83 ± 9.66%) exceed the across session chance level (8.28 ± 0.85%; paired t test, t(4) = 3.36, p = 0.028, n = 4 mice, Cohen’s d = 2.19; Fig. 5c-f). The assembly patterns from post-task REM sleep on the first day were present in the pre-task REM (mean = 96.30 ± 3.70%, across session mean = 11.76 ± 3.40%; paired t test, t(4) = 16.82, p = 0.00007, Cohen’s d = 10.96, n = 4), suggesting that there is preserved assembly structure from naïve REM in both the task and later REM. We then identified assembly patterns in REM sleep recorded before the first track exposure. Each pre-task assembly was subsequently reactivated at least once during exploration of the novel track (Fig. 5e), indicating that assemblies present in the naïve state are recruited by later experience, with some of these assemblies containing member cells with similar future place fields (Fig. 5d). However, naïve pre-task assemblies were detectable in wake at a similar rate when templates were applied to other subjects and sessions (paired t test - t(4) = 2.15, p = 0.099 | Cohen d = 1.40).

Next, we applied sequence factorization to REM sleep recorded before the novel experience and identified consistent, reliable ordered sequences composed of distinct, non-overlapping neuronal populations (Fig. 5g,h). Extending this analysis across all combinations of pre-task REM, post-task REM, and wakefulness (Fig. 5i), we found no difference in the number of sequences between the preplay condition (12.75 ± 1.79 events; 0.04 ± 0.004 Hz) and the replay condition (9.75 ± 1.79 events; paired t test, t(3) = 1.029, p = 0.38, n = 4 mice; Fig. 5i). Thus, ordered sequences were already present in REM sleep before the corresponding experience, and at a rate comparable to post-experience replay.

## Discussion

Hippocampal reactivation has been examined almost exclusively within the framework of NREM sleep, in which sharp-wave ripples compress place-cell sequences into brief, temporally coordinated events^4,5,7^. REM sleep, by contrast, is a state for which evidence of structured reactivation has been sparse and inconsistent, described in early reports as either absent or only occasional^14,15^, and in recent large-scale recordings, as weak reactivation during REM-associated theta^17^. In this context, a notable and unanticipated finding of the present study is that REM sleep is instead dominated by the robust re-expression of co-active cell assemblies. Because assemblies were defined during awake locomotion and subsequently reactivated during REM, both theta-dominated states^13^, this mode of reactivation appears to operate independently of the ripple-based framework.

Assembly reactivation was experience-dependent and most pronounced under emotionally salient conditions. Following exposure to an anxiogenic environment, assemblies reactivated more frequently and preferentially recruited their most strongly contributing neurons, whereas sequential replay was unchanged in both frequency and the fidelity of individual events. REM sleep has been implicated in the consolidation of emotionally salient memory^19,27–29^, and its theta rhythm is causally required for contextual fear memory^11^; however, the network-level operation underlying this role has remained undefined. The reactivation of adult-born dentate granule cells during REM sleep supports fear-memory consolidation^18^, but does not specify the form of the reactivated activity.

Our results identify a potential mechanism: rather than simply rehearsing behavioral trajectories, REM sleep directly modulates the participation of core assembly members. Anxiogenic experience increases the participation of these core cells, potentially strengthening the neural representation of salient experiences, but this enhanced participation diminishes as the aversive environment becomes familiar. This dynamic modulation suggests that REM sleep assembly reactivation preferentially reinforces salient experiences while adapting to their changing behavioral relevance.

Novelty engaged a distinct, and in several respects opposite, operation. A single novel exploration increased the frequency of ordered sequential replay, to our knowledge the first demonstration that experience selectively up-regulates ordered sequence replay within REM sleep, while the overall rate of assembly reactivation remained unchanged. Assembly composition was nonetheless refined, with weakly contributing neurons pruned from subsequent reactivations and strongly contributing members retained. This process parallels, at the level of network membership, the synaptic homeostasis proposed for REM sleep, whereby weakly potentiated connections are eliminated and those engaged during learning are preserved^35^. Novelty and anxiety thus gave rise to a double dissociation: novelty enhanced sequential replay and refined assemblies through subtraction, whereas anxiety amplified assembly reactivation through recruitment and left sequential replay unaltered. This dissociation aligns with the roles proposed for each form of reactivation, in which ordered sequences represent the relational structure of a continuous experience and co-active assemblies mediate the associative binding of discrete, salient events^36,37^.

A further level of organization was present prior to the relevant experience. In REM sleep preceding an animal’s initial encounter with the track, we recovered both co-active assemblies and ordered sequences that were subsequently expressed during exploration, a sign of preconfigured internal dynamics^24^ rather than a residual trace of the preceding waking state. These sequences relate to preplay as described during NREM sleep^30–32^, with one methodological distinction: NREM preplay was defined by Bayesian decoding of spatial trajectories, whereas the present events are defined by preserved temporal order without positional reference, the level of structure accessible in our data. We therefore adopt “preconfigured” as the umbrella term and reserve “preplay” for its ordered form. Although the existence of preplay remains debated^33^, the finding that hippocampal dynamics are shaped by developmental factors independent of experience^34^ lends it a plausible substrate. Preconfigured sequences were expressed at rates comparable to post-experience replay, indicating that novel experience does not generate sequential activity de novo. Replay was nevertheless more frequent after novel than familiar exposure, suggesting that familiarity suppresses sequence expression. Finally, experience reorganized the network: assembly membership was pruned after novelty and reinforced after anxiogenic exposure, and exploration activity correlated more strongly with the subsequent than the preceding REM episode. Experience thus appears to select and refine a pre-existing scaffold rather than generate structure anew.

Several aspects of our approach may account for why these events have largely eluded detection. The principal earlier demonstration of temporally structured replay in REM sleep relied on template matching in animals already well habituated to their environment^15^, while subsequent studies sampled modest numbers of neurons or applied a single detection method^17^, conditions poorly suited to the sparse firing of REM and to reactivation that is most robust following novelty. By combining large populations with complementary methods spanning progressively stronger assumptions about temporal order, co-activity^25,38,39^, Bayesian decoding^40,41^, and sequence factorization^26^, and by cross-validating against conservative surrogates, we recovered structure that any single approach could fail to detect. This layering proved essential: Bayesian decoding of spatial trajectories alone yielded no above-chance replay, despite the clear presence of ordered sequences recovered by factorization. Because sequence factorization identifies recurring temporal firing motifs without reference to the animal’s position, we characterize these events by their occurrence and their preserved neuronal order rather than by any inferred spatial trajectory, a distinction that also cautions against interpreting them as literal representations of movement through space.

The implications of these findings extend beyond the hippocampus. REM sleep declines with normal aging, is disproportionately disrupted in Alzheimer’s disease^42–44^, and is dysregulated in anxiety- and stress-related disorders, particularly PTSD^45–48^, precisely the conditions under which memory consolidation and emotional regulation deteriorate. By resolving distinct REM operations for novel and for aversive experience, our findings offer candidate mechanisms through which disrupted REM sleep could compromise the consolidation of recent experience and the regulation of emotional memory. Central questions for future work are how REM and NREM reactivation are coordinated across sleep, and whether the assembly- and sequence-based operations described here are selectively vulnerable in disease.

## Methods

### Animals

Parvalbumin-Cre mice (The Jackson Laboratory, stock number 017320) were used under protocol (DOUG-5206/7650) and guidelines approved by McGill University and the Canadian Council of Animal Care. Mice were housed in individual cages with 12:12 light cycles. A total of 2 female and 5 male mice were used. In the anxiety task, one mouse did not sufficiently explore the open arm to reliably quantify activity there and was therefore excluded for that task.

### Surgery procedure

All surgeries were conducted under deep anesthesia with isoflurane. AAV5-CaMKII-GCaMP6f (AAV5.CamKII.GCaMP6f.WPRE.SV40, Penn Vector Core via Addgene) was diluted by 1:2 and 200 nL was injected at dorsal CA1 region (ML: 1.5, AP: -2.0, Depth: -1.3) using a Nanoject III at 1nL/s speed (Drummond). Two weeks after the injection, a gradient refractive index (GRIN) lens (diameter 1.8 mm) was implanted over the injection site. Tungsten microelectrodes were inserted into the contralateral CA1 region (ML: -1.5, AP: -2.0, Depth: -1.3). Reference and ground electrodes were inserted in the frontal lobe and cerebellum. The entire implant was firmly secured with anchored screws and dental cement (C&B-Metabond, Patterson Dental Canada). Four weeks after implantation, a baseplate for the miniscope was chronically fixed in the region of interest and a flexible wire was additionally inserted into the neck musculature for EMG recording. All the microelectrodes and wires were soldered to an 8-channel electric interface board (EIB-8, Neuralynx) and connected to the Neuralynx system via an HS-8 preamplifier (Neuralynx, USA).

### Behavioral tasks

Following surgery, mice were individually housed in the sleep-recording room for one week before recordings began. Handling and habituation with the recording equipment were performed daily by the same experimenter. The task session (10 min) was recorded around 12 pm after recording pre-task sleep. Mice were water-deprived one day before the start of the experiment, and their weight was monitored daily. During the task, mice explored a linear track (1 m long, 10 cm wide, and 80 cm high, with 8.5 cm high walls), and water containing 10 % of sucrose was delivered at each end of the track after each run completed by the mouse. Since mice were never exposed to the task before the start of the experiment, day 1 was considered a novel experience, and day 5 was considered a familiar experience. The position of the mouse on the linear track was tracked using an LED light on the headstage of the miniscope. To introduce spatially-confined anxiety on the linear track, the same elevated linear track was modified to have walls removed along half the track length. Mice were trained to collect rewards on this anxiogenic linear track for 6 days.

### Electrophysiology and calcium imaging

All the recordings were performed during the light cycle in order to maximize the number of sleep episodes. Neural activity during sleep was recorded in the home cage before (pre-task sleep) and after the task (post-task sleep). Each sleep recording session lasted ∼2 h. During each sleep session, CA1 LFP were recorded along with EMG signals and behavioral movements. Using these signals, REM sleep states were manually classified by the experimenter in real-time based on the presence of strong consistent theta oscillations and a flat EMG signal. When the LFP and EMG met these criteria the experimenter would start the calcium recording. To prevent photobleaching of the calcium reporter, the maximum duration for calcium imaging was limited to a total of 30 min per day. Additional offline classification was performed using the ratio of theta/delta band power and root mean square of the EMG to restrict recordings to REM periods. All calcium imaging recordings were acquired at 30 frames per second.

### Calcium imaging processing

For a given mouse and recording day, all calcium imaging videos from sleep and wakefulness were concatenated into a single video, while retaining the associated timestamps. Motion was corrected between each frame using NoRMCorre^49^. Subsequently, calcium transients for each neuron were extracted using constrained non-negative matrix factorization (CNMFe)^23^. We used the following parameters for the calcium extraction: gSig=3, gSiz=15, background_model=’ring’, min_corr=0.6, min_PNR=8. For further analysis, calcium imaging datasets were analyzed using Python 3.10.0 and MATLAB (Mathworks, Natick, MA).

### Spatial tuning

Calcium transients were binarized as described previously^40^. Briefly, animal positions were discretized in 2.5 cm bins. For each neuron and spatial bin, the activity likelihood, the marginal likelihood (regardless of location), and the prior probabilities (occupancy rates) were extracted. To estimate the spatial tuning strength, we used an adjusted version of mutual information that scales between 0 (random tuning) and 1 (perfect association) and penalizes low sampling using mutual information computed on a null model distribution^40^. Additionally, the significance of this association between neural activity and spatial locations was assessed using a Chi^2^ test of independence. Among neurons exhibiting a significant association between position and neural activity, neurons were ranked according to their adjusted mutual information, and the top *n* = 256 neurons were selected for subsequent analyses. This number was chosen based on the most conservative estimate of the number of sequences detected during sequential analysis and was kept consistent across all the analyses. No filtering was applied to the resulting tuning curves so as to prevent spurious spatial relationships in downstream replay analyses.

### Unstructured assembly detection

Ensembles of co-active neurons were established as described previously^25,38,39^ with minor adjustments to apply these methods to calcium data. Briefly, a correlation matrix was generated using the z-scored binarized activity of the top 256 spatially tuned neurons in 0.5s time bins (Figure S1). For awake data, only the activity during periods of mobility (>9cm/s) was included.

The eigenvalues of the correlation matrix which passed the lambda-max threshold of a Marčenko-Pastur distribution were considered ‘candidate assemblies’ (Figure S1c-d). Independent component analysis (ICA) using the ‘FastICA’ package (500 shuffles) was applied to the z-scored data using the number of candidate assemblies to obtain a weight score for the contribution of each neuron to the candidate assemblies (Figure S1e). The assemblies whose ICA weights did not include at least 3 neurons with strong positive weights (>1.96 SD) were excluded. The normalized weights of each candidate assembly were then applied to z-scored and binned (0.5s) binarized data to determine the expression strength of that assembly in time (Fig. S1f-j). To determine the chance level for assembly expressions, we circularly shifted the activity of each neuron by a random number of samples to create a surrogate data set, computed the expression strength based on the original ICA weights, and repeated this process 500 times. Assemblies expression was considered significant for bins where the projection crossed the 99th percentile of the shuffled expression strength (mean REM ‘R threshold’: 4.71±1.06). The across session responses, described below, were used to z-score the within session metrics (Fig 5e-g). For Figures 2-4 assembly templates were extracted using the wake data and applied to REM data, while Fig 5 the pre-REM data was used to extract assembly templates which were then applied to wake data.

As an alternative to within-session shuffle surrogates to determine the chance levels of assembly activation/reactivation, we also applied the templates to the data within the same session (‘within’) and data from all other sessions (‘across’). This allowed us to test chance detection using intact hippocampal data. To determine if assembly templates from one of the three states (pre-REM, wake, and post-REM), we applied templates (reference) against data across all sessions (target), subsampling the target data to match the size of the reference data when needed. To measure the relative changes in assembly membership, the participation of each cell was assessed by taking the probability of being active in each assembly reactivation and normalizing it by the out of assembly firing probability (Fig 3c, d, Fig S1n). To determine if experience altered the participation of cells, we took the participation probability in the post-task REM and divided it by the normalized pre-task probability to give a participation change score.

### Spatial Bayesian replay

To evaluate the occurrence of Bayesian replays (i.e. neural activity representing trajectories with varying speed), we leveraged the probabilistic tuning curves and marginal likelihood values computed above, as well as binarized activity during sleep, to compute posterior probabilities as described previously^40^ (Figure S2). Importantly, we used a uniform prior to ensure that the distribution of previous explorations did not bias the representations of locations during offline periods. Posterior probabilities were smoothed temporally by computing the product between posteriors of two adjacent frames (covering ∼ 66 ms). Using these posterior probabilities, we then computed the maximum a posteriori (MAP), expressed as putative locations being replayed in offline periods. We then evaluated the null hypothesis that replayed locations are distributed randomly. To this end, we computed a linear fit on 0.5 s windows for actual neural data and shuffled surrogates, where the spatial and temporal structures are randomly shuffled using circular permutations. On both actual and shuffled datasets, we then computed a goodness-of-fit score (R^2^) as well as the maximum spatial interval between two adjacent frames (jumpiness), based on an approach described previously^41^. For each temporal window, a significant replay event was considered only if the score for those two metrics outperformed shuffled surrogates in 95 % of cases (n = 1,000 surrogates). Since mice displayed a bias towards exploring the closed arm more than the open one in the anxiety task, we controlled for this potential confound by subdividing the linear track in 4 equal parts, and sub-sampling exploration using the smallest number of samples along the track. Then, using this balanced sampling approach, tuning curves and Bayesian probabilities were established as described above.

### Ordered sequences

Finally, to monitor the occurrence of ordered hippocampal sequences, we employed non-negative matrix factorization of binarized neural data. Specifically, we implemented seqNMF^26^. Briefly, we expressed our neural data matrix *X* as a product of *n* sequences *W* by time *H*:

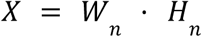

To evaluate sequence replay, we computed W during the awake exploration of the linear track and multiplied that matrix with that of neural activity during subsequent REM sleep:

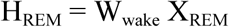

When sequences were established and tested in the same recording, we used a 50/50 train/test ratio. To evaluate the significance of ordered sequences, we also computed *H* using matrix multiplication between sequence patterns where the identity of neurons is scrambled. This process was repeated for n = 1,000 shuffled surrogates, and only times where *H_actual_*exceeded *H_shuffled_* more than 95% of the time were sequence occurrences considered significant. SeqNMF *lambda* and *L* parameters were optimized over a range of values (Fig. S3). Sequence scores were directly derived from actual and shuffled *H* values using:

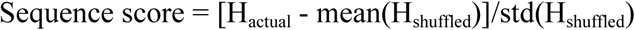

### Histology

After completing experiments, mice were anesthetized with a ketamine/xylazine/acepromazine (3 mg/kg) through intraperitoneal (IP) injection. Mice were perfused transcardially with 4% paraformaldehyde (PFA) with PBS. Brains were fixed with PFA for another day at 4 °C and the solution was changed to PBS for the next day. Brains were sliced with 50 µm thickness using a vibratome. The location of recording electrodes, GRIN lenses, as well as the expression of GCaMP6f in the pyramidal layer of CA1 was confirmed with post-mortem histological analyses.

### Statistics

Unless noted otherwise, all data are reported as mean ± SEM. Parametric tests were used whenever data was normally distributed and variance homogeneous, otherwise non-parametric tests were employed (and described appropriately). RM-ANOVA, repeated-measures ANOVA; *, p < 0.05; **, p < 0.01; ***, p < 0.001; ****, p < 0.0001; ns, not significant.

## Acknowledgments

We would like to thank Javad Karimi Abadchi for valuable feedback on the manuscript as well as the members of the Williams and Brandon labs for helpful comments. This work was supported by funding from the Canadian Institutes for Health Research (CIHR) Foundation Program FDN-148478, CIHR Project grants 191680 and 186113, the Natural Sciences and Engineering Research Council of Canada (NSERC) Discovery Grant RGPIN-2020-06717, and a Tier 1 Canada Research Chair to S.W. GE was supported by Brain Canada, the Roger J. Paiement Outreach Award and the Douglas Institute Foundation. JC was supported by a Healthy Brain Healthy Lives (HBHL) fellowship, Gerald Clavet Fellowship, and Hugh E. Burke Fellowship. JEC was supported by an Alzheimer’s Association and by the Canada Brain Research Fund (AARF-22-928742), an innovative arrangement between the Government of Canada (through Health Canada), the Brain Canada Foundation, and the Alzheimer’s Association, and a Conrad F. Harrington Fellowship. The funders had no role in study design, data collection and analysis, decision to publish, or preparation of the paper.

## Code availability

All codes and software used to generate the results of this study are publicly available on Github:

- All source codes used in the current study are available along with instructions here: https://github.com/Jisoochoi1127/Replay_REM)
- Tuning curves are extracted using: https://github.com/etterguillaume/PyCaAn (versions 0.1.1)
- Extraction of calcium imaging data was done using: https://github.com/etterguillaume/MiniscopeAnalysis (version 1.0)
- This extraction pipeline leverages motion correction from NoRMCorre: https://github.com/flatironinstitute/NoRMCorre (v0.1.1)
- This pipeline leverages CNMFe video processing for calcium traces extraction: https://github.com/zhoupc/CNMF\_E (v1.1.2).

## Dataset availability

All data used to generate the results of this study will be made available publicly in a repository upon publication. All data is available early to reviewers upon request to the authors.

## Contributions

Conceptualization: GE, JC, SW

Methodology: GE, JC, JEC

Formal Analysis: GE, JEC, JC

Investigation: JC

Visualization: JEC, JC, GE

Funding acquisition: SW, JC, JEC, GE

Project administration: SW Supervision: GE, SW

Writing – original draft: JC, GE, JEC

Writing – review and editing: JC, JEC, GE, SW

---

**Supplemental figure S1.**
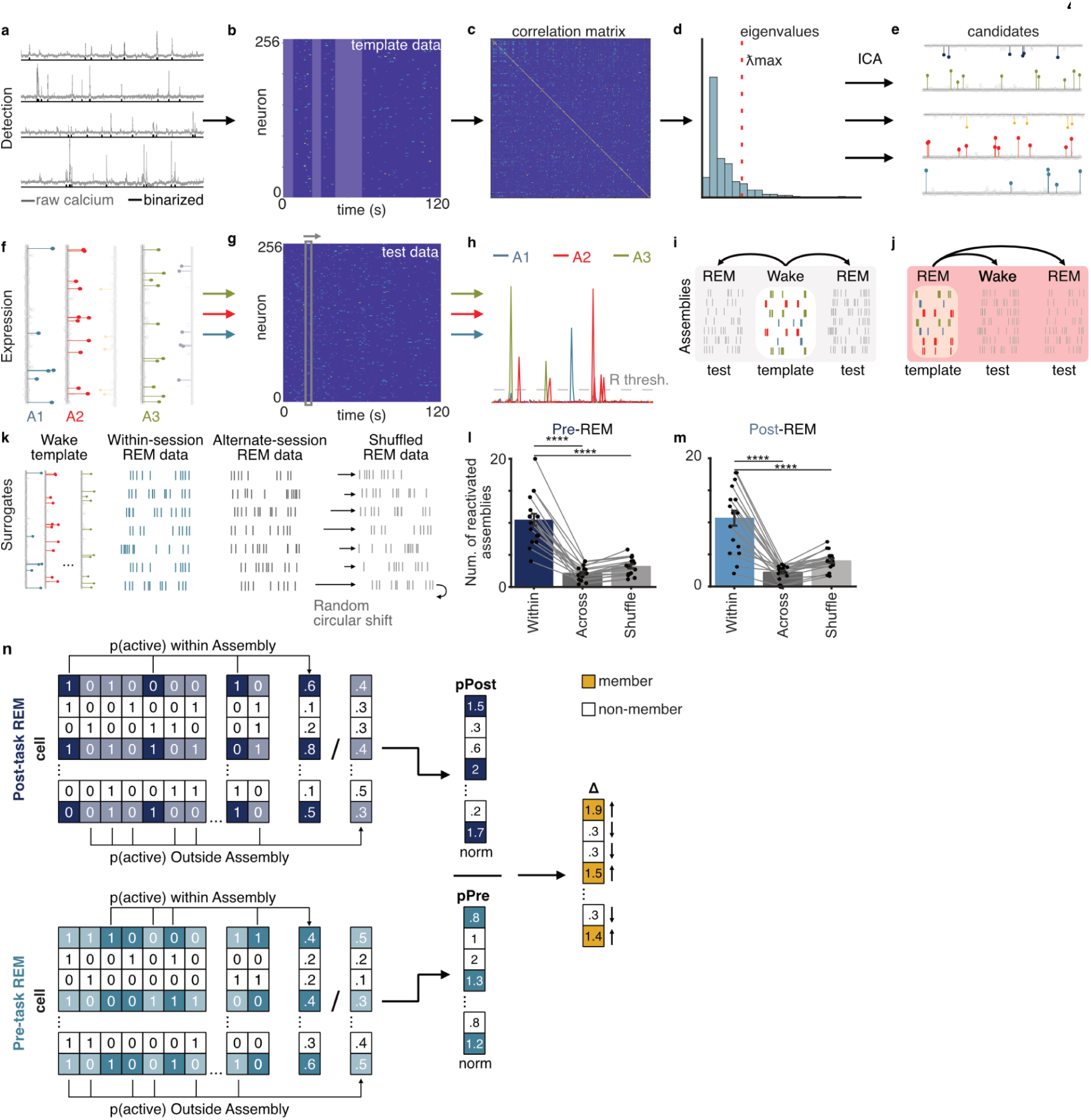
Assembly detection and expression process. **a**, Example calcium traces with binarized activity (black) from the top 256 place cells. **b**, Z-scored activity with periods of low mobility (<9cm/s; opaque) removed. **c**, Activity correlation matrix. **d**, Histogram of eigenvalues. The lambda-max value from the Marčenko-Pastur distribution is used as a threshold to determine the number of highly-correlated assemblies to be extracted using ICA. **e**, Examples of candidate assemblies from ICA with weights for each neuron. **f**, Assemblies with positive weights are kept and applied to the z-scored activity of a test data set (**g**) to determine the expression strength of each assembly (**h**). Shuffled data is used to determine the significant reactivation threshold (’R thresh’) and any assemblies that do not show significantly strong expression compared to shuffled surrogates are removed. **i,** To determine if awake co-activity patterns are present in REM sleep, wake data is used as the template and REM is used as the test. **j**, Pre-REM patterns are also extracted and applied to subsequent wake and post-REM data for Figure 5. **k**, To assess the chance level of reactivations in REM data, wake templates were applied to both pre- and post-REM data within a session and against two surrogate sets: those from alternate sessions (mean counts from all other subjects/sessions), and from the 500 shuffles of the within session REM data. Within session REM data contained significantly more assemblies passing inclusion criteria (**h**) than surrogates (F(2, 53) = 43.14, p < 0.0001) using pre-REM data (**l**). **m,** Tukey-Kramer post-hoc comparisons revealed a significant difference between ‘within’ (10.22 +/- 1.12) and ‘across’ (2.21 +/- 0.22) conditions (p<0.0001) and between ‘within’ and ‘shuffle’ (3.89 +/- 0.32; p<0.0001), but not between the surrogate conditions (p=0.1680). **n**, Visualization of the assembly participation metric. The activity of each cell across post-task REM assembly reactivations, normalized by the out of assembly activity, is divided by the normalized pre-task REM activity to yield the change in participation for member (shaded) and non-member (unshaded) cells. Data expressed as mean ± SEM. ****, *p* < 0.0001. Test used in **l/m**: repeated measures ANOVA.

**Supplemental figure S2.**
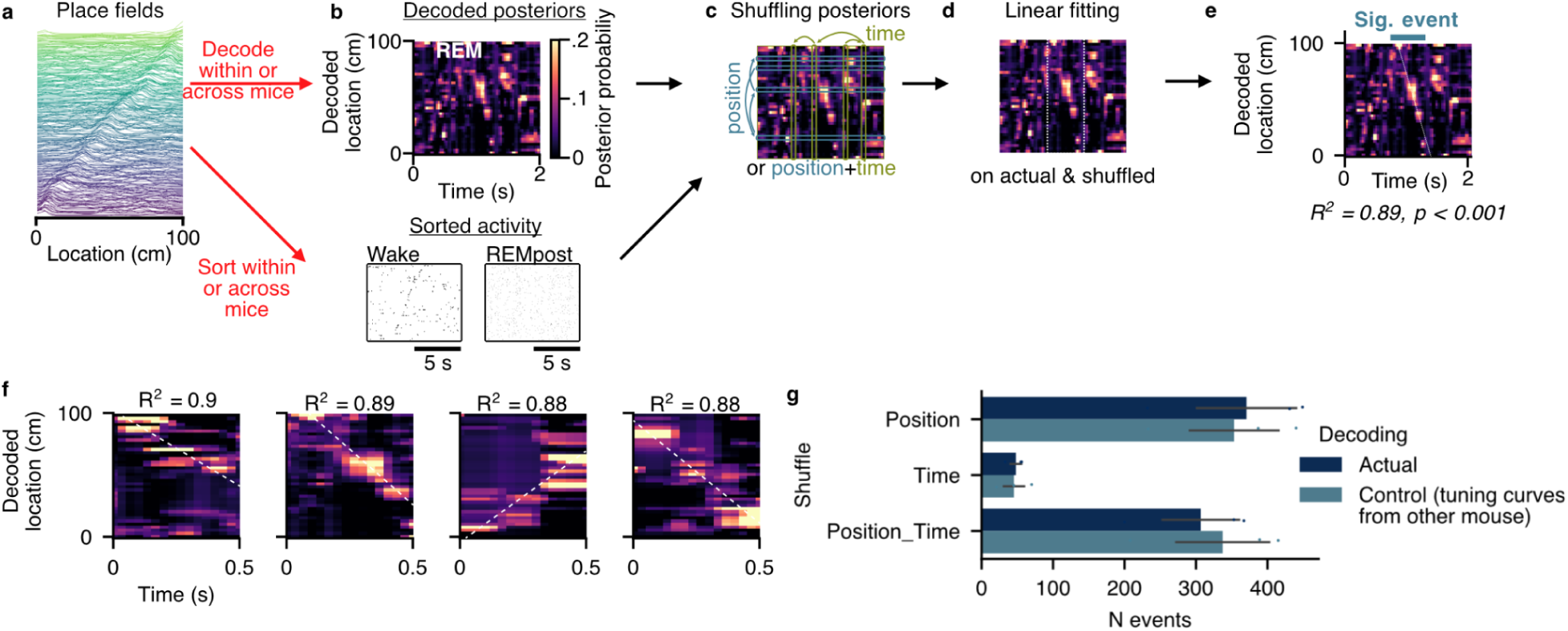
Bayesian-based approach for detecting replay of trajectory. **a**, For each neuron, spatial information is computed and place fields are extracted based on spatial information. **b**, Using the known tuning curves of each neuron (or those of a randomly selected mouse, to compute ‘across subject’ controls, used as a control to estimate false-discovery rate), Bayesian posterior probabilities are computed during REM (pre and post; top). Alternatively, raw binary activity is sorted using the peak location of each place field (2.5cm bins). **c**, For both approaches, posterior probabilities (or sorted binary activity) are shuffled in the time, position, or both dimensions at the same time (n= 1,000 surrogates). **d**, Replayed locations are estimated using maximum a posteriori (MAP) or average activity index (for raw binary data). A line-fitting procedure was then applied using a 0.5-s sliding window to both the actual and shuffled data. **e**, Goodness-of-fit (R^2^), jumpiness, and replay length are computed for actual data and shuffled surrogates, for each non-empty window where jumpiness is non-zero (i.e. more than a single location is represented by the putative replay event). From these putative replay events, only those with a R^2^ value exceeding 95% of the shuffled surrogates are considered to be significant events. To apply a Bonferroni correction, the 0.05 probability threshold (alpha level) is divided by the number of windows used to test for significance. The same procedure is applied for sorted binary activity within, and across mice. **f**, Examples of linear replay candidate events. **g**, Cross-validation of Bayesian sequences within and across sessions. For across session validation, the tuning from the reference session was applied to the REM data from another session. Data expressed as mean ± SEM. ns, not significant. Test used in **f/g**: two-way ANOVA.

**Supplemental figure S3.**
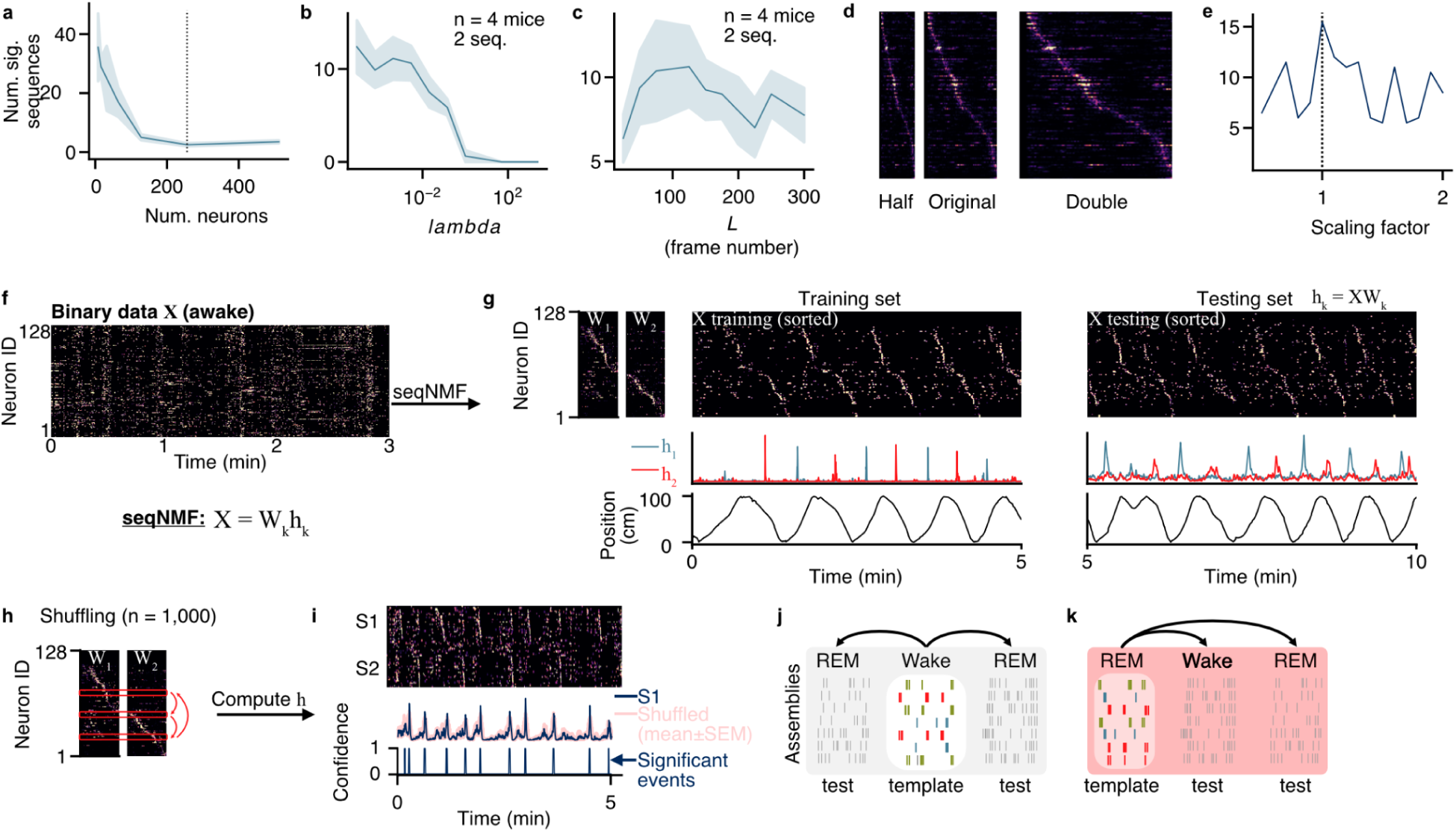
Optimization of SeqNMF. **a**, Number of significant sequences, assessed against chance, across different neuronal sample sizes. The sample size used in this study (256 neurons) yielded the most conservative estimate of the number of sequences detected during REM sleep. **b**, Number of significant sequences replayed during REM as a function of lambda. **c**, Same but as a function of L, the window size used to establish sequence templates. **d**, Example sequence template (center) scaled with factors 0.5 (left) and 2 (right). **e**, Number of significant sequences replayed during REM sleep following awake experience after scaling sequence templates. **f**, SeqNMF assumes that raw data X can be factorized into k sequences (W_k_) and temporal vector (sequence strength; h_k_) using X = W_k_h_k_. Here, an example recording during wakefulness is used to extract a template but REM sleep preceding experience can also be used to this end. **g**, Sequence templates W, sequence strength h, and raw data sorted using sequence templates for the first (training set) and second half of linear track exploration. **h**, To test for significance, the identities of neurons within sequence templates are shuffled (n = 1,000 surrogates). Sequence strength h is computed using actual data and shuffled surrogates. **i**, On a testing set, significant replay events are defined as epochs for which sequence strength exceeds that computed using shuffled surrogates 95% of the time. **j**, To assess replay, awake experience is used as a training set, and sequences are monitored in REM preceding (pre) and succeeding (post) this awake experience. **k**, To evaluate pre-configured sequences and pre-play, REM-pre is used to extract putative sequences, and sequence significance is evaluated during wakefulness. Data expressed as mean ± SEM.

**Supplemental figure S4.**
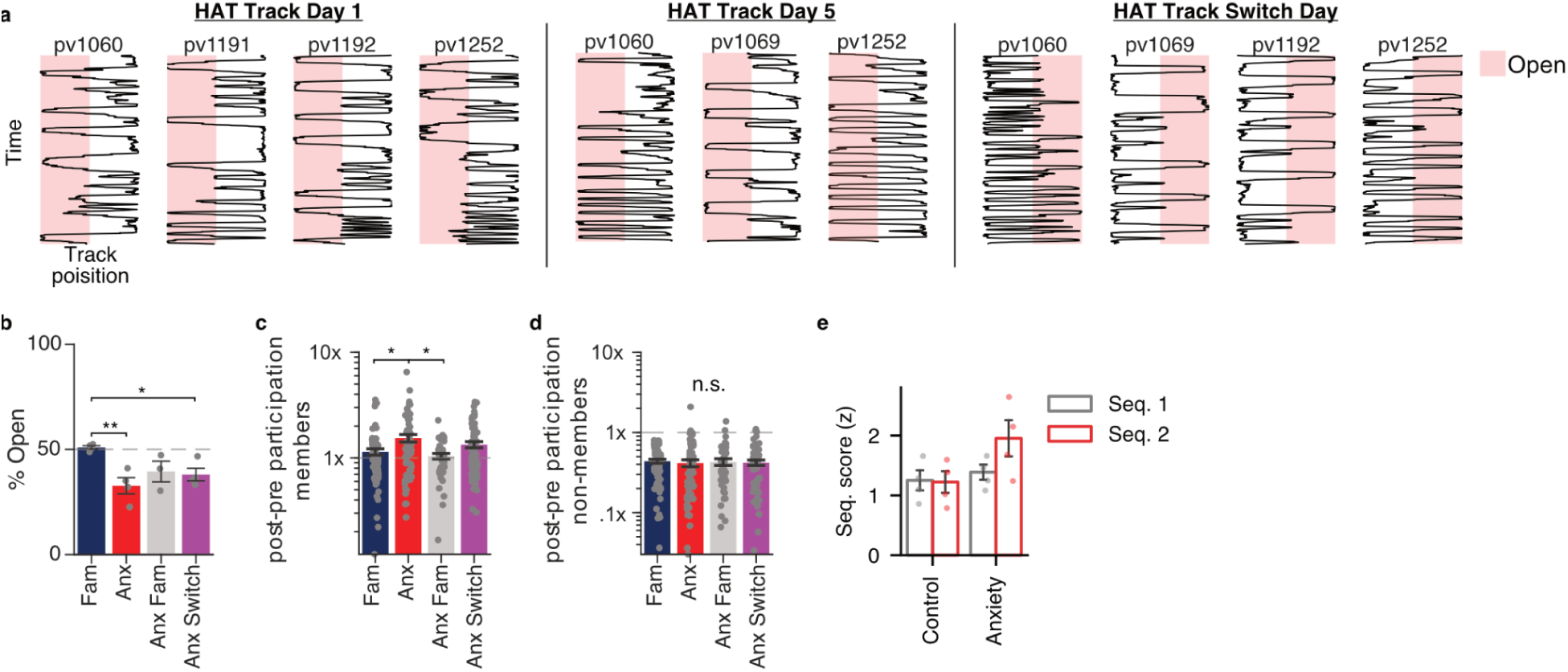
Behavioural on the anxiety task. **a**, Panels represent anxiety task sessions. Day 1 was the first exposure to the half-open track. Day 5 represents a familiar condition on the track followed by a reversal of the open section on the Switch day. The x axis shows mouse location on the 100cm linear track, and the y axis time (10min session). **b**, The track day significantly impacted the time spent in the open arm of the track (1-way ANOVA, F(3,9): 7.81, p < 0.01) with Day 1 ‘Anx’ (32.72 +/- 3.84%, p < 0.01) and ‘Switch’ (38.04 +/- 2.98%, p = 0.04) showing reduced open-arm occupancy relative to the fully enclosed familiar session (50.93 +/- 0.81%; Tukey-kramer post hoc comparisons). **c**, Track conditions impacted the participation of assembly member cells (one-way ANOVA, F(3,244): 4.83 p = 0.003) with Day 1 (1.54 +/- 0.13) showing a significant enhancement compared to the enclosed familiar (1.14 +/-0.08, p = 0.013, tukey-kramer corrected) and anxiety familiar sessions (1.04 +/- 0.06, p = 0.005, tukey-kramer corrected). **d**, The participation of non-member cells did not change across conditions (one-way ANOVA, F(3,245): 0.10 p = 0.96). **e**, Sequence scores did not differ between anxiety track and familar controls. Data expressed as mean ± SEM. ns, not significant; *, *p* < 0.05; **, *p* < 0.01. **b-d**: 1-way anova with Tukey-kramer post hoc comparisons.

